# Focused ultrasound increases gene delivery to deep brain structure following the administration of a recombinant adeno-associated virus in the cerebrospinal fluid

**DOI:** 10.1101/2024.02.09.579587

**Authors:** Rikke Hahn Kofoed, Kate Noseworthy, Kathleen Wu, Laura Marie Vecchio, Chinaza Lilian Dibia, Shuruthisai Sivadas, Sheng-Kai Wu, Kristina Mikloska, Malik White, Bradford Elmer, Shyam Ramachandran, Christian Mueller, Kullervo Hynynen, Isabelle Aubert

## Abstract

Gene delivery via adeno-associated viral vectors can provide lasting clinical benefits following a one-time treatment. Delivery throughout the brain is needed for the treatment of neurological disorders with widespread pathology, including Alzheimer and Parkinson diseases, and amyotrophic lateral sclerosis. Most gene vectors have poor diffusion in the brain tissue. Furthermore, it is only at high intravenous doses that gene vectors can overcome the blood-brain barrier. In contrast, relatively lower doses of gene vectors injected in the cerebrospinal fluid enable significant transduction of superficial brain regions. The remaining challenge and unmet need of gene therapy is to deliver gene vectors to deep brain structures using a minimally invasive strategy. Here, we demonstrate that non-invasive focused ultrasound blood-brain barrier modulation can increase the delivery of recombinant adeno-associated virus by 5-fold to deep brain structures following injection in the cisterna magna. Delivery of adeno-associated viral vectors to the central nervous system, via administration in the cerebrospinal fluid, is being evaluated in several clinical trials for treating beta-galactosidase-1 deficiency, Batten disease, Alzheimer disease, Parkinson disease, amyotrophic lateral sclerosis, and spinal muscular atrophy. Our findings suggest that the efficacy of gene therapies delivered in the cerebrospinal fluid can be enhanced by targeting brain areas of interest with focused ultrasound.

**Significance statement:** Administration of viral vectors in the cerebrospinal fluid through the cisterna magna is being evaluated in patients with neurological disorders. Focused ultrasound combined with intravenous microbubbles safely increases the permeability of the blood-brain barrier in humans and enables delivery of intravenous adeno-associated virus in non-human primates. Here, we demonstrate that combining these two clinically relevant gene delivery methods, intracisterna magna administration and focused ultrasound with microbubbles, can facilitate gene delivery to superficial and deep brain structures. Our findings have the potential to increase the efficacy of gene therapies, particularly for disorders with brain regions that have remained difficult to reach.

## Introduction

Gene therapy has demonstrated long-lasting therapeutic benefits for the treatment of neurological disorders affecting large parts of the central nervous system, such as spinal muscular atrophy(1). Recombinant adeno-associated virus (AAV) is the most advanced vector for gene delivery *in vivo*, and some AAV serotypes, such as AAV9, can cross the blood-brain barrier (BBB) after intravenous administration(2). However, BBB crossing of AAV9 requires high intravenous dosages of up to 2×10^14^ genome copies per kilogram (GC/kg) in children and more in adults where the ability of AAV9 to cross the BBB is limited(2, 3). The recent deaths of patients with X-linked myotubular myopathy receiving 3×10^14^ GC/kg AAV8 highlight the risks associated with intravenous AAV administration and the need to develop new strategies for AAV delivery to the central nervous system(4). The permeability of the BBB can be increased transiently by the application of focused ultrasound combined with intravenous microbubbles (FUS-MB)(5). FUS induces an oscillation of the microbubbles, which decreases tight-junction proteins and increases transcytosis across the endothelial cells(6, 7). The temporary increase in BBB permeability facilitates non-invasive delivery of intravenous AAV to FUS-targeted brain areas at dosages 50-100 times lower than needed for BBB crossing by AAV9 alone(8). Still, the delivery is limited to FUS-targeted brain areas, and while FUS can be targeted simultaneously to multiple brain regions the current FUS methods remain unsuitable for treating whole-brain diseases(9).

The first gene therapies tested in Alzheimer and Parkinson diseases were delivered by intracranial injections of AAV which resulted in persistent transgene expression for over 10 years but limited to the injection site(10, 11). Injection of AAVs in the cerebrospinal fluid (CSF) through the cisterna magna is gaining momentum in clinical trials focusing on whole-brain diseases, because of the resulting AAV distribution to several brain areas while being relatively less invasive than intra-cerebroventricular (ICV) delivery; however, the gene delivery following intracisterna magna administration is limited to superficial brain areas(12, 13). We hypothesized that combining FUS-MB with intra-CSF AAV administration would expand the transduction profile of AAV to include FUS-targeted deep brain structures (Figure 1A), enabling translation of this technology for the treatment of neurodegenerative diseases where both superficial and deep brain structures are affected.

**Figure 1.**
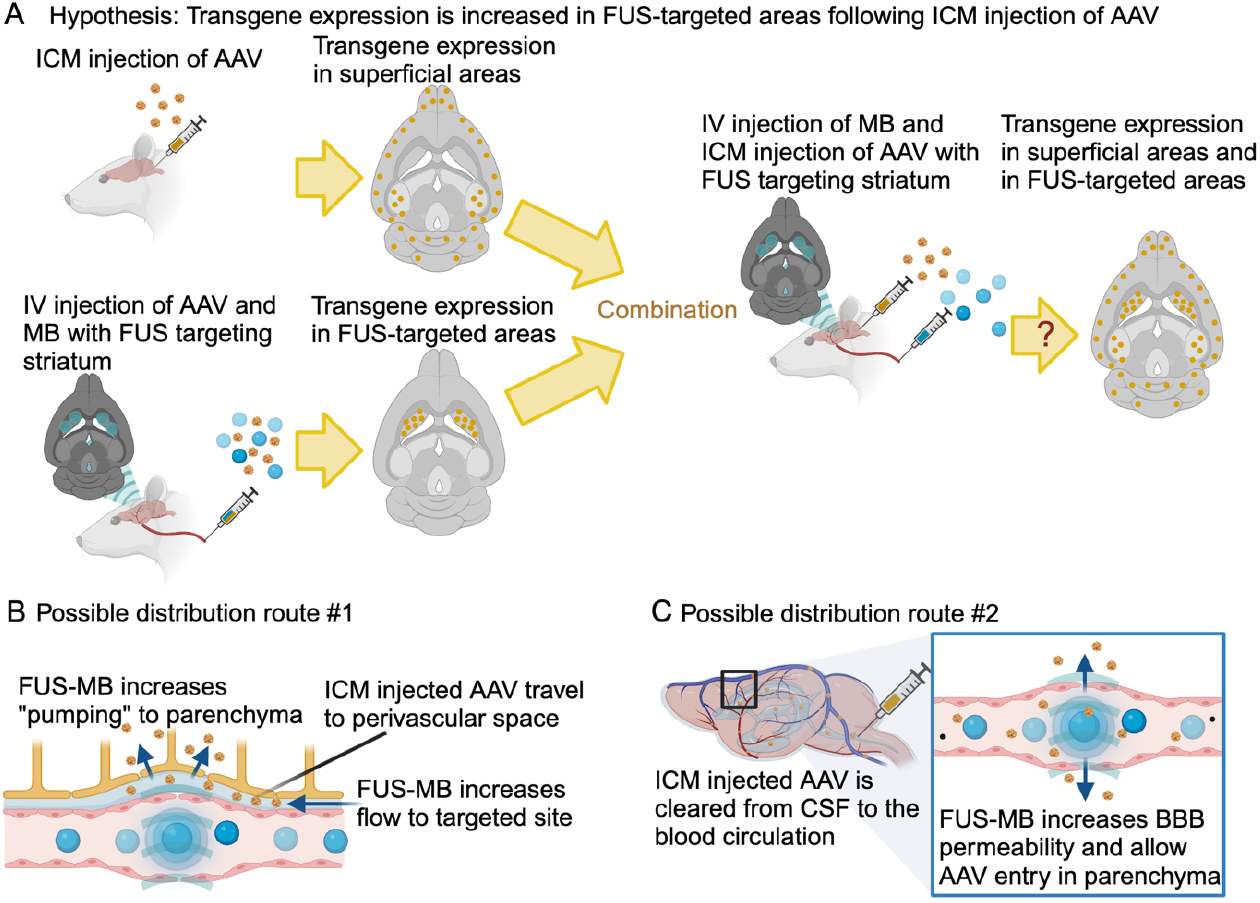
Hypothesis and possible mechanisms of action. **A** Intracisterna magna (ICM) administration of AAV leads to widespread transduction, especially of superficial brain areas, but the transduction of deeper brain structures (e.g. striatum) is limited. Focused ultrasound (FUS) combined with intravenous (IV) microbubbles and AAV can deliver genes to any FUS-targeted brain region, including deep structures; however, the transduction is only seen in the FUS spots. We hypothesize that combining FUS and IV microbubbles with AAV administered ICM can lead to both gene delivery widespread to superficial brain areas and to deep brain structures targeted with FUS. **B, C** There are at least two possible mechanisms of action describing how FUS and microbubbles can increase brain delivery of ICM-administered AAV. **B** Possible distribution route #1 hypothesizes the AAV travels in the CSF to the perivascular space, and from there, the interaction between FUS and IV microbubbles creates a pumping effect, which increases the distribution of the AAV from the CSF into the brain parenchyma. **C** Possible distribution route #2 is based on the fact that ICM-injected AAV will eventually get cleared from the CSF into the blood. Then, the AAV could enter the brain from the blood at FUS-targeted sites, similarly to when the AAV is injected intravenously(16).

At least two potential mechanisms of action could mediate FUS-MB-induced brain delivery of ICM-injected AAV: First, AAV could travel in the cerebrospinal fluid to the perivascular space, where the interaction between FUS and microbubbles can induce a pumping effect increasing the flow of AAV to the perivascular space in the FUS-targeted area and into the brain (Figure 1B). This distribution route was previously suggested for FUS-MB brain delivery of intranasally-administered AAV(14). The second possible mechanism is based on the fact that AAV injected in the CSF eventually distributes into the blood(15). It is possible that AAV travels from the cisterna magna into the blood and then enters the brain at FUS-MB targeted sites where the BBB has increased permeability (Figure 1C). The two proposed mechanisms are not mutually exclusive.

The results presented in this study demonstrate poor transduction of the striatum by AAV after ICM administration, and that FUS-MB increases delivery to this brain region without the need for peripheral dosing of vector. Our data suggests that the mechanism of action is principally related to the proposed distribution route #2 (i.e. FUS-MB-mediated brain delivery of AAV that has distributed from the CSF to the blood). Distribution route #1 (i.e. FUS-MB-mediated perivascular pumping) could not be proven or ruled out with the current data.

## Results

### Kinetics of AAV distribution following ICM delivery and timing relative to FUS-MB application

To elucidate the ability of FUS-MB to increase brain delivery of ICM-administered AAV, we performed a series of pilot studies to determine the optimal timepoint for AAV injection relative to FUS-MB. Distribution route #1 (Figure 1B) depends on the AAV being present in the perivascular space at the time of FUS-MB application to take advantage of a putative pumping effect. To determine the kinetics of substances injected ICM in our rat model we administered an MR contrast agent, gadolinium, ICM and performed MR imaging 45, 55, 70 and 80 minutes (min) post administration (Figure 2A). Gadolinium enhancement (white) was clearly seen in brain areas proximal to the subarachnoid space containing cerebrospinal fluid; e.g., cortical areas, hippocampal formation, cerebellum, and ventral part of midbrain and brainstem (Figure 2B-E). Gadolinium enhancement increased with time in brain structures further away from the subarachnoid space, such as the thalamus, from 45 min to 80 min post administration (gadolinium enhancement (white) increased in Figure 2e1 compared to Figure 2b1). This suggests a gradual diffusion of the ICM-injected gadolinium into the brain parenchyma, perhaps via the perivascular space. The striatum remained without significant gadolinium enhancement at the time points investigated, and it was chosen for FUS-targeting as a brain region with poor access to ICM-injected substances in this rat model. Because the diffusion of gadolinium into the brain parenchyma continued from 45 min to 80 min post-injection and AAV is larger (>3,500 kDa) than gadolinium (∼0.5 kDa), 120 min was chosen as a possible timepoint when ICM-injected AAV could be present in the perivascular space. It remained possible that small amounts of gadolinium in the perivascular space have not been visible on the MR images. Therefore, to avoid the potential of losing a significant amount of AAV from the perivascular space into the blood before FUS-MB application, 60 min was also chosen as a timepoint for FUS-MB application following AAV injection ICM. A pilot study was conducted to determine the feasibility of FUS-MB delivery of ICM-injected AAV to the brain using five treatment groups. Group 1: as a positive control of FUS-MB brain delivery, animals were injected intravenously with AAV during FUS-MB application as routinely performed (Figure 2F)(9, 16, 17). Groups 2 and 3: AAV was injected 120 min and 60 min, respectively, before FUS-MB application to determine FUS-MB delivery to the brain of ICM-injected AAV (Figure 1G). Group 4: to determine the relevance of distribution route #2 (Figure 1C), a group of animals was injected with AAV 10 min post-FUS-MB application to investigate if AAV is delivered to the FUS-targeted site without potential perivascular pumping (i.e. no AAV in the perivascular space at the time of FUS-MB application) (Figure 1G). Group 5: a group of animals was injected with AAV ICM without FUS-MB application (Figure 1H). FUS was targeted bilaterally in two spots in the striatum (Figure 1F+G). We utilized the modified AAV2, AAV2-HBKO, which provides increased distribution in the brain parenchyma following intravenous injection combined with FUS-MB targeting the striatum, as well as a decreased uptake in the liver compared to AAV9(16). AAV2-HBKO encoded green fluorescent protein (GFP) under control of the ubiquitous promoter CAG. AAV2-HBKO was administered at a dose of 2.16×10^12^ GC/kg irrespective of the administration route, which is within the range previously used for FUS-MB-mediated AAV delivery to the brain following intravenous injection (1.67×10^12^ GC/kg – 1.67×10^13^ GC/kg)(16). Four weeks following AAV delivery and bilateral FUS-MB targeting to the striatum, the animals were sacrificed and tissues were harvested for subsequent analyses. One hemisphere was used for immunohistochemical (IHC) analysis of protein expression, and the other hemisphere was used for reverse transcription quantitative polymerase chain reaction (qPCR) analysis of mRNA expression (Figure 2J). Images of sections stained with anti-GFP antibodies showed GFP expression in the FUS spots of all FUS-MB treated groups, irrespective of route of administration and injection timepoint, though with varying levels of expression (Figure 2K). In animals injected intravenously with AAV, GFP expression was only visible in the FUS spots, whereas in animals injected ICM with AAV, expression was visible in FUS spots as well as in cortical regions, hippocampus, and the midbrain (Figure 2K). Analysis by qPCR of the FUS-targeted striatum showed the highest GFP mRNA levels in animals injected intravenously with AAV during FUS-MB, followed by animals injected ICM with AAV 120 min prior to FUS-MB treatment (Figure 2L). Animals injected ICM with AAV 60 min prior to FUS-MB and 10 min post FUS-MB showed similar levels of GFP mRNA expression, trending towards a higher level of expression than animals injected ICM with AAV without FUS-MB (Figure 2L). It was not possible to conduct a meaningful statistical analysis of the pilot study due to the variability between animals and the low number of animals per group (n=3-4). However, the IHC analysis confirms GFP expression in FUS-targeted spots and hence the ability of FUS-MB to modulate brain delivery of AAV injected ICM. In addition, the observed GFP protein expression in the FUS spot in animals injected ICM with AAV 10 min post-FUS-MB suggests that distribution route #2 is possible because in this group the AAV is not in the perivascular space when FUS-MB and the potential pumping effect is applied.

**Figure 2.**
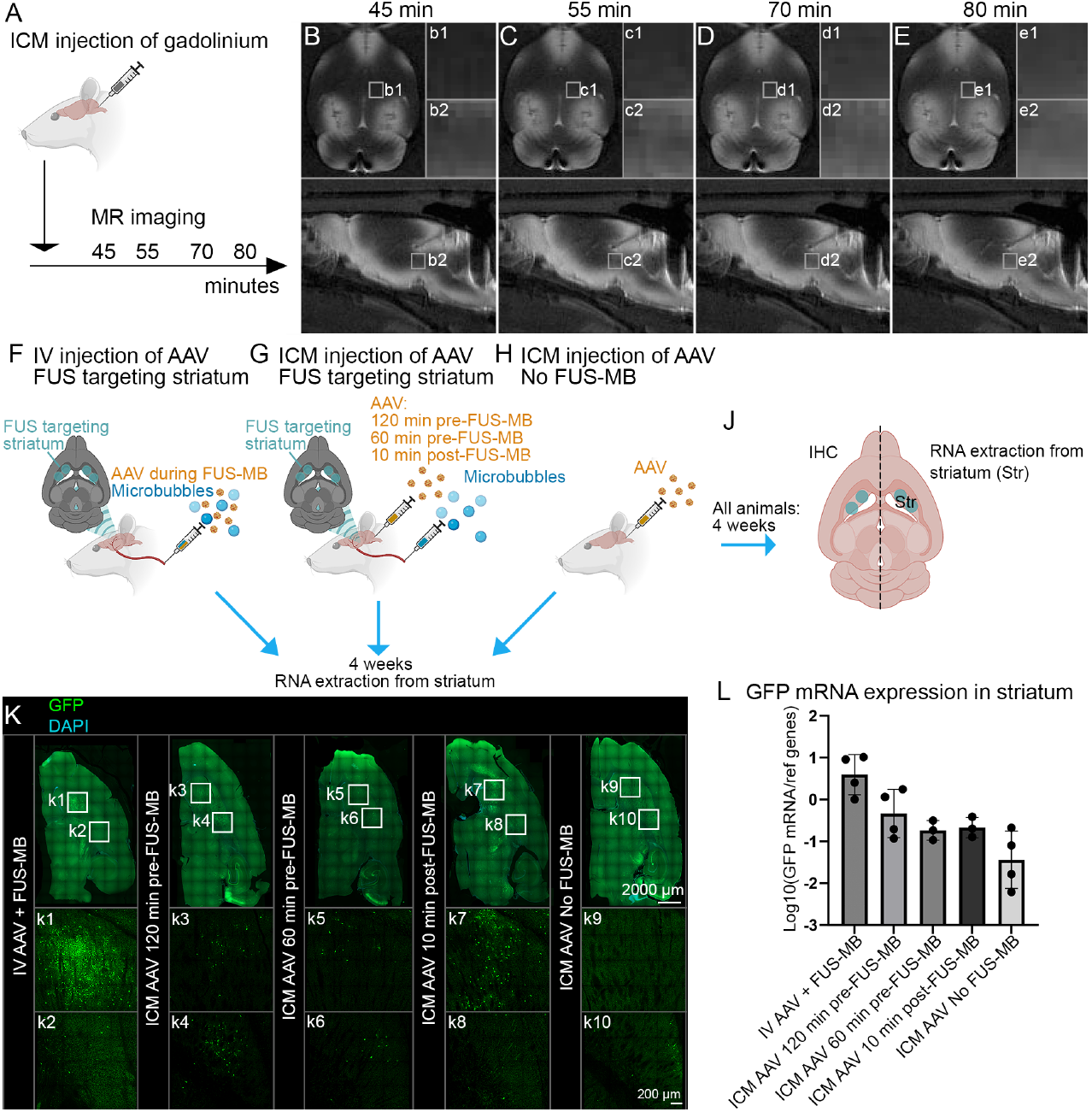
Pilot trial of AAV dosing timepoints relative to FUS-MB. **A** To determine the diffusion of ICM-injected particles into the brain parenchyma (some of which could be via the perivascular space), the MR contrast agent gadolinium was injected ICM and **B-E** T1-weighted MR images were acquired after 45, 55, 70, and 80 min. **F-H** Five groups were tested for FUS-MB-mediated delivery of AAV: **F** AAV was injected intravenously during FUS-MB application as a positive control, **G** AAV was injected ICM either 60 min or 120 min prior to FUS-MB application or 10 min post-FUS-MB application. The injection of AAV after FUS-MB application was used to evaluate whether distribution route #2 was possible. **H** One group of animals received ICM injection without FUS-MB application. **J** After 4 weeks, the animals were sacrificed, and one hemisphere was used for GFP immunohistochemistry (IHC) and the other for GFP mRNA expression via qPCR. **K** GFP protein expression was seen in the FUS-targeted spots in all animals except those not treated with FUS-MB, where GFP expression was visible in only one area of the striatum close to the cortex (k10). **L** Pilot study with small group sizes (n=3-4 per group) showed GFP mRNA expression in the FUS-targeted striatum with the highest expression in animals injected IV with AAV during FUS-MB followed by AAV injection ICM 120 min before FUS-MB. Injection of AAV ICM 60 min before FUS-MB and 10 min post-FUS-MB resulted in GFP mRNA levels trending towards a higher gene delivery than ICM injection without FUS-MB.

### ICM-injected AAV is significantly cleared into the blood

To further investigate the influence of distribution route #2 (i.e. FUS-MB-mediated brain delivery of AAV that has distributed from the CSF to the blood), we compared the levels of AAV that are cleared from the CSF into the blood and reaching peripheral organs following ICM administration and direct AAV intravenous injection. To that effect, viral vector genome copies were quantified in peripheral organs four weeks following AAV administration (Figure 3A). There were no significant differences between genome copies in the spleen, kidney, muscle, liver, heart, and lung measured by digital droplet PCR (ddPCR against GFP) analysis in animals injected intravenously with AAV compared to animals injected ICM (Figure 3B). This suggests that most AAV-injected ICM was cleared from the cerebrospinal fluid into the blood to transduce peripheral organs. This data also supports the hypothesis that, at a given time point post-ICM administration, the AAV concentration in the blood becomes high enough to lead to a significant crossing from the blood into the brain at FUS-MB-targeted sites following ICM injection of AAV (Figure 1C). To determine the level of AAV in the blood at the time points for FUS-MB treatment following intravenous (during FUS-MB) and ICM (60 min and 120 min before FUS-MB) injection of AAV, blood samples were obtained as soon as possible following FUS-MB treatments, resulting in blood samples from 60 min and 153 min post-ICM injection and 21 min post intravenous injection (Figure 3C). Analysis of GFP genome copies (i.e., AAV) in the blood samples demonstrated a tendency toward increased AAV levels 60 min after ICM injection. However, this was not significantly different from negative control samples from animals not injected with AAV with the current sample size of n = 3-6 (Figure 3D). Blood samples taken 153 min post ICM injection demonstrated a significant increase in GFP genome copies compared to 60 min after injection (Figure 3D). The level of AAV in the blood 21 min post intravenous administration was approximately twice the level of AAV 153 min post ICM injection (Figure 3D). Previous studies in mice have demonstrated that doubling the intravenous dosages of AAV2-HBKO do not significantly affect the percentage of transgene-positive cells obtained in the brain following FUS-MB delivery(16). Because of the significant clearance of AAV from the cerebrospinal fluid into the blood after 153 min, distribution route #2 (Figure 1C) is likely to be the primary delivery mechanism responsible for GFP protein and mRNA expression seen in animals injected ICM with AAV 120 min prior to FUS-MB (Figure 2K+L).

**Figure 3.**
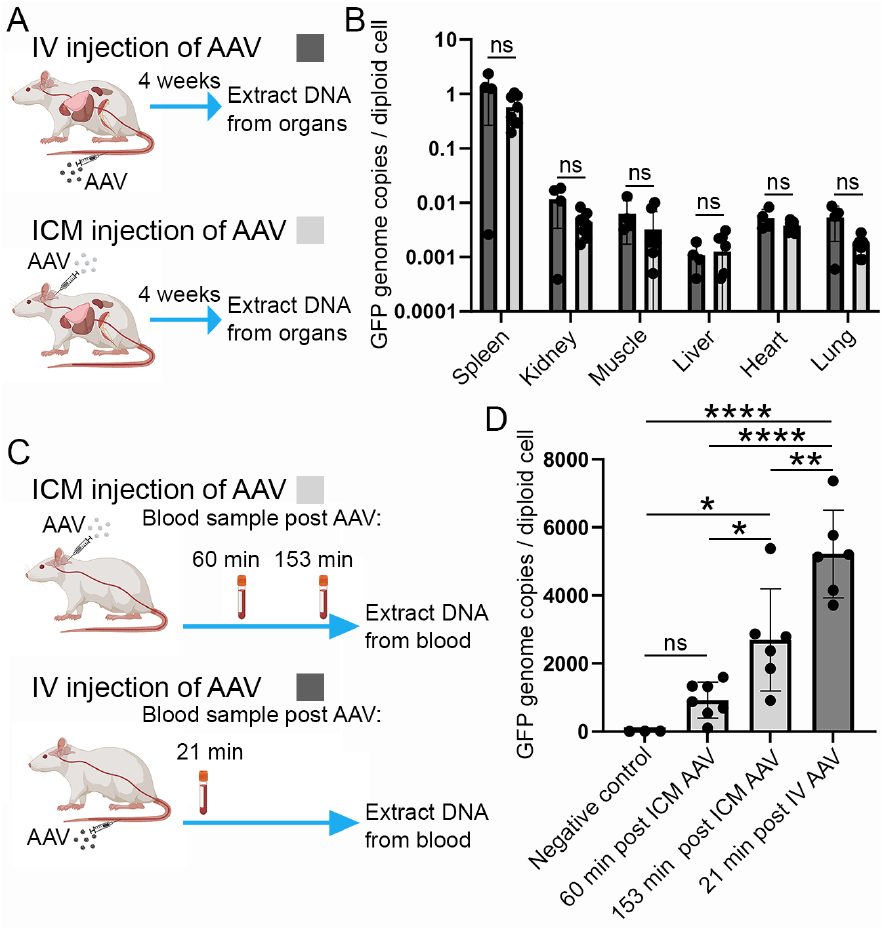
AAV is significantly cleared from the cerebrospinal fluid into the blood. **A** The distribution of AAV to peripheral organs following intravenous (IV) and ICM administration was measured by quantifying the GFP genome copies in the peripheral organs four weeks after AAV administration. **B** There were no significant differences between the GFP genome copies in the peripheral organs with IV injection of the AAV compared to ICM injection. **C** To determine the kinetics of AAV distribution in the blood following IV and ICM injection, GFP genome copies were quantified in blood samples taken 60 and 153 min post ICM injection and 21 min post IV injection. **D** GFP genome copies in the blood 60 min after ICM injection was not significantly higher than negative control samples. GFP genome copies in the blood 153 min after ICM injection was significantly higher than the 60 min time point and the negative control. The level of GFP genome copies was the highest of all groups 21 min after IV injection of the AAV. Bars represent mean +/- standard deviation. Statistical analysis was done using one-way ANOVA and post hoc Tukey’s test, ****p < 0.0001, **p < 0.01, *p < 0.05, n = 4-6/group (n = 3 for negative control in D).

### FUS-MB increases delivery of ICM-injected AAV to the striatum

The pilot study demonstrated a tendency towards higher GFP mRNA expression in the striatum when AAV was injected ICM 120 min prior to FUS-MB compared to 60 min prior to and 10 min following FUS-MB. In addition, the blood samples suggest a higher level of intravenous AAV 120 min compared to 60 min following ICM administration. In a larger cohort of animals, AAV delivery was therefore compared between animals injected intravenously with AAV during FUS-MB, animals injected ICM with AAV 120 min prior to FUS-MB, and animals injected ICM with AAV without FUS-MB application (Figure 4A). Animals were sacrificed after four weeks and brains analyzed by IHC and qPCR (Figure 4A). GFP protein expression (green, white arrows) in the brain were seen in FUS-targeted brain areas in animals injected both intravenously and ICM with AAV, but not in animals injected ICM without FUS-MB application (Figure 4B). Analysis of GFP mRNA expression in the FUS-targeted striatum showed increased levels of GFP mRNA expression in animals injected with AAV intravenously and ICM and treated with FUS-MB compared to animals injected ICM with AAV without FUS-MB application (Figure 4C). There was no significant difference in GFP mRNA expression in the striatum in animals injected with AAV intravenously and ICM and treated with FUS-MB (Figure 4C). The data in Figure 4C was log10-transformed to obtain normal distribution, and the non-transformed data showed a 5.6-fold increase in GFP mRNA expression in the striatum following FUS-MB application in animals injected ICM with AAV compared to animals without FUS-MB treatment (Supplemental Figure 1). GFP mRNA levels were also measured in brain areas not targeted with FUS-MB, where gadolinium enhancement in the pilot study suggested a significant delivery of ICM-injected particles (Figure 2B-E). ICM administration of AAV, with or without FUS-MB targeting the striatum, resulted in significantly increased levels of GFP mRNA expression in the thalamus, midbrain, cerebellum, hippocampus, and brainstem compared to animals injected intravenously with AAV (Figure 4D-H). The depth of the FUS spots in the z-axis also targets brain structures located dorsally and ventrally relative to the striatum (Figure 4A, last diagram to the right, turquoise ovals). There were no significant differences between groups in the cortical structures (Cortex 1 and 2; CX1 and CX2) that FUS spots partially covered (Figure 4I+J). However, in the cortical area Cortex 3 (CX3), which was not affected by FUS-MB, there was a significantly higher level of GFP mRNA in animals injected ICM with AAV compared to intravenously (Figure 4K). The data presented in Figure 4 demonstrate that FUS-MB can increase delivery to deep brain structures, such as the striatum, even with ICM-injected AAV, which does not otherwise reach these structures efficiently. Importantly, ICM injection of AAV leads to significantly increased gene delivery to brain areas that are not targeted by FUS-MB compared to intravenous AAV.

**Figure 4.**
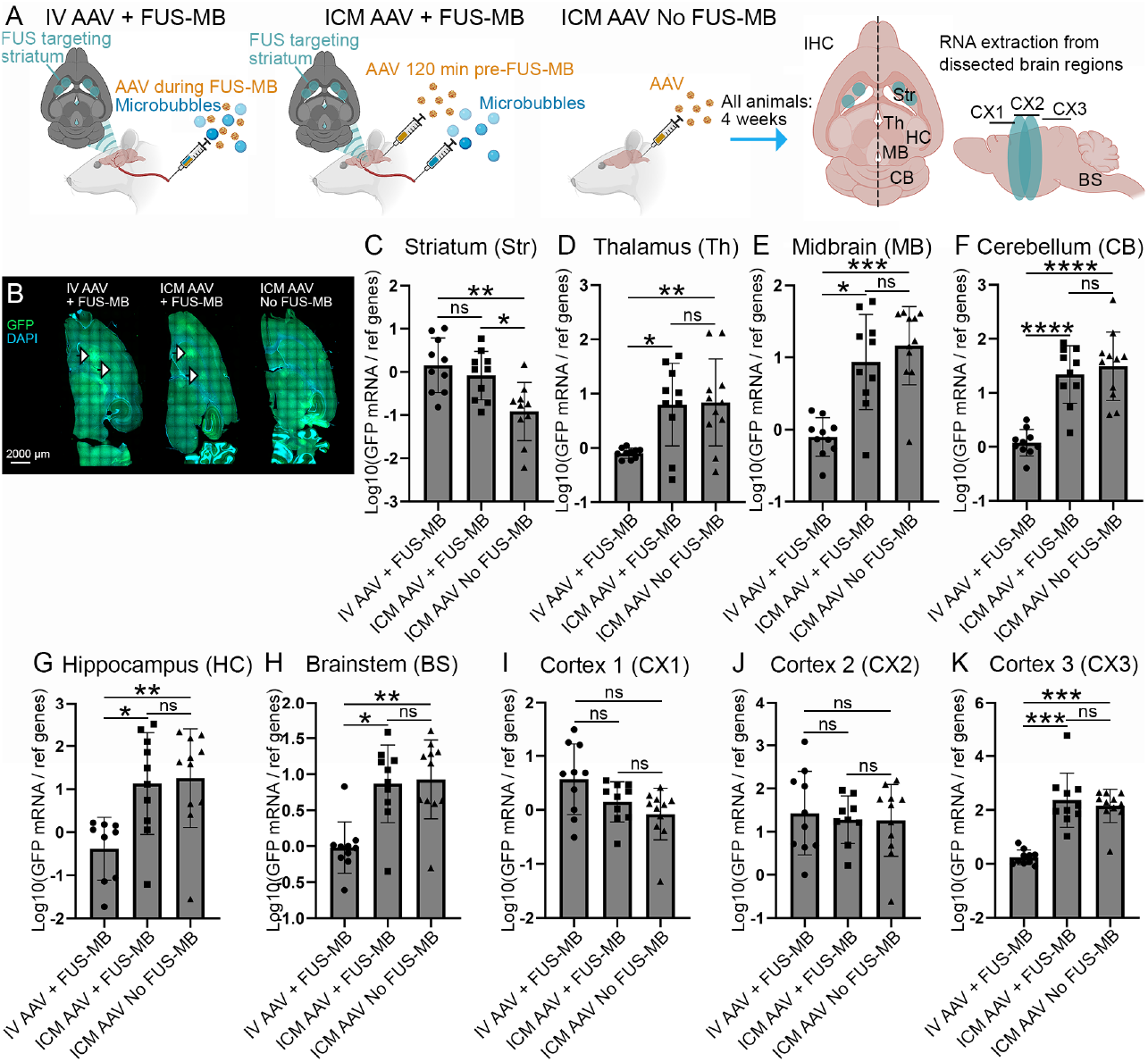
FUS-mediated delivery of ICM administered AAV to the striatum. **A** AAV was injected intravenously (IV) during FUS-MB, ICM 120 min pre-FUS-MB, or ICM without FUS-MB application. FUS was targeted bilaterally to the striatum. After 4 weeks, the animals were sacrificed, one hemisphere was used for IHC and the other was dissected into brain regions for RNA extraction and qPCR analysis. **B** IHC staining showed GFP expression in the FUS-targeted spots in the striatum (green, white arrows) in both animals injected with AAV IV and ICM, but not in animals injected with AAV ICM without FUS-MB application. **C-K** GFP mRNA expression was quantified in multiple brain regions. **C** In the striatum, IV AAV + FUS-MB, and ICM AAV + FUS-MB resulted in significantly higher GFP mRNA expression than ICM AAV without FUS-MB. There was no significant difference in GFP expression in the striatum between animals injected with AAV IV and ICM and treated with FUS-MB. **D-H** ICM injection of AAV, both with and without FUS targeting the striatum, resulted in significantly higher GFP mRNA expression than IV injection in the thalamus, midbrain, cerebellum, hippocampus, and brainstem. **I-J** When targeting the striatum, the FUS also hits cortical structures (see A). AAV ICM injection did not result in higher GFP mRNA expression in the FUS-targeted cortical structures compared to animals injected with AAV IV and treated with FUS-MB. **K** In cortical structures not hit by FUS, there was significantly higher GFP mRNA expression in animals injected ICM with AAV, both with and without FUS-MB application in the striatum, than in animals injected IV with AAV and FUS-MB. Data was not normally distributed and was therefore log10-transformed, which provided normal distribution for the data from the striatum, thalamus, cerebellum, and cortex 3. Log10-transformation did not achieve normal distribution for data from the hippocampus, brainstem, midbrain, cortex1, and cortex2. These brain areas were statistically analyzed using the non-parametric Kruskal-Wallis test and post hoc Dunn’s test. Statistical analysis of all other brain areas was done using one-way ANOVA and post hoc Tukey’s test i, ****p < 0.0001, ***p < 0.001, **p < 0.01, *p < 0.05, n= 10-11/group. Non-transformed data can be found in Supplemental Figure 1.

## Discussion

### FUS-MB increases gene delivery to deep brain structures following ICM administration

The field of gene therapy is rapidly evolving. Monogenic disorders with a known underlying genetic cause are receiving particular attention with a high potential to be treated with gene therapy(12). The first monogenic disease that was treated with a one-time gene therapy treatment was spinal muscular atrophy using an intravenous administration of AAV9 in a dose of 1.1×10^14^ GC/kg (final recommended dose)(1). The ability of AAV9 to cross the BBB and blood-spinal cord barrier in infants ensures that the therapy can reach target cells in the central nervous system. However, high intravenous doses come with the risk of severe side effects and the decreased permeability of the BBB with age means that higher dosages are required to treat diseases that occur later in life(2, 4). Direct delivery of AAV in the CSF is emerging as a promising route of delivery to facilitate wide-spread gene delivery to the brain and spinal cord with higher efficiency than when using intravenous delivery(12, 15, 19–21). While administration in the CSF through the cisterna magna results in gene delivery to multiple brain regions, delivery to deep brain structures, such as the striatum, remains limited(15).

Here, we demonstrate that FUS-MB can increase the delivery of ICM-administered AAV2-HBKO to the striatum in rats. The delivery to the FUS-targeted striatum with ICM and IV AAV administration was not significantly different; however, the ICM route resulted in higher gene delivery to non-FUS-targeted areas than IV injection. Taken together, our results suggest that FUS-MB can be used to increase the transduction volume following ICM administration of AAV to include FUS-targeted deep brain structures, which cannot be reached efficiently by ICM injection alone. Increased delivery to deep brain structures by FUS-MB can enhance the therapeutic efficacy of gene therapy treatments targeting diseases affecting the entire brain and ensure a better distribution of gene delivery across the brain.

### Mechanism of action

There are at least two possible routes of distribution that can explain the increase of ICM AAV delivery to the brain with FUS-MB (Figure 1B+C). The first possible route is based on an emerging hypothesis proposed for FUS-MB-mediated gene delivery following intranasal administration of AAV (Figure 1B). Intranasally administered AAV is thought to move into the CSF and from there to the perivascular space, where the interaction between FUS and intravenous microbubbles results in a pumping effect, which increases the movement of AAV from the perivascular space to the brain parenchyma(14). The same group also showed increased distribution of ICM-administered fluorescently-labeled albumin to the perivascular space following FUS-MB application compared to without FUS-MB(22). Whether FUS-MB increases distribution of particles to the perivascular space, induce a pumping effect that helps particles to enter the parenchyma, or both of these mechanisms, remain to be determined. In the current study, three major differences are observed compared to previous studies on FUS-MB-mediated brain delivery of intranasally administered AAV and dextrans: 1) Intranasal administration of AAV5 results in significantly lower levels of transgene genome copies in peripheral organs compared to intravenous AAV5 injection(14). Using AAV2-HBKO in rats, we observed no significant difference in transgene genome copies in peripheral organs between animals injected intravenously and ICM. 2) FUS-MB application results in higher gene delivery to the brain following intranasal injection of AAV5 compared to intravenous injection(14). This may partly be due to the increased retention of AAV5 in peripheral organs following intravenous administration(9). We did not observe significant differences in gene delivery to the FUS-targeted brain area with FUS-MB following intravenous and ICM administration of AAV2-HBKO. Despite the use of different AAV serotypes, points 1) and 2) together suggest a different distribution route of AAV to the FUS-targeted area and the periphery following intranasal administration compared to ICM administration. 3) A study combining intranasal delivery of dextrans with FUS-MB showed increased distribution to FUS-targeted brain areas when FUS-MB was applied after, and not before, intranasal administration(23). Though only a small number of animals were tested (n=3), we observed transgene expression in the FUS-targeted brain area when FUS-MB was applied 10 min before ICM injection of AAV2-HBKO. Overall, there are apparent differences in the mechanism of action when using FUS-MB to increase brain delivery following intranasal administration of particles compared to ICM. However, because FUS-MB affects the diffusion of albumin to the perivascular space following ICM administration, it remains possible that FUS-MB also enhances the distribution of ICM-injected AAV to the perivascular space(22).

The hypothesized distribution route #2 (Figure 1C) is based on the fact that AAV administered into the CSF is eventually cleared from the CSF into the periphery, which we also confirmed in this study(15). Therefore, AAV may be transported from the CSF into the blood and from here to the brain parenchyma at FUS-targeted sites with increased BBB permeability. Two findings in the current study suggest that this route plays a significant role in FUS-MB-mediated delivery of ICM-injected AAV to the brain: 1) Transgene expression was observed in the brain when FUS-MB was applied before ICM injection of AAV. The pumping effect described in distribution route #1 depends on particles being present in the perivascular space during FUS-MB application. Therefore, distribution route #1 is not possible when FUS-MB is applied before AAV administration, suggesting a different mechanism of action is responsible. With the currently available data by us and others, however, it is impossible to determine the relative impact of a potential increase in the distribution of ICM-injected particles to the perivascular space following FUS-MB treatment(22). 2) There was no significant difference in GFP mRNA expression in the FUS-targeted striatum between animals injected intravenously and ICM with AAV. In addition, the level of AAV in the blood shortly after FUS-MB application (measured 21 min post intravenous administration and 153 min post-ICM administration) only shows a two-fold difference in animals injected intravenously and ICM. Because a two-fold increase in intravenous AAV2-HBKO dosages did not previously result in significant differences in brain transduction, the AAV blood concentration 120 min after ICM administration is high enough to explain why similar levels of transduction are seen in the FUS-targeted striatum following intravenous and ICM injection of AAV(16).

### Translation and future perspectives

With the currently available data, further investigations of ICM + FUS-MB as a delivery strategy of AAV to the brain should be designed to account for both possible distribution routes #1 and #2. It is impossible to completely exclude either pathway for FUS-MB-mediated delivery of AAV-administered ICM. Considerations should include the timepoint of FUS-MB application following ICM administration and uptake of the gene vector in peripheral organs. A high uptake of gene vectors in peripheral organs can limit the amount in free circulation in the blood and, as such, their FUS-MB delivery to the brain(9).

Future investigations should focus on further investigating the mechanisms of action, which are essential to optimize the design of large animal studies and, ultimately, clinical translation. While the pumping effect is difficult to image in real-time, the distribution of AAV in the perivascular space at different time points following ICM administration and FUS-MB application could be imaged with high-resolution imaging or electron microscopy(22). In addition, we have previously demonstrated that AAV5 is only delivered to a very low extent to the brain following intravenous administration because of the high peripheral uptake(9). This feature of AAV5 may be useful to determine the significance of distribution route #2 in FUS-MB delivery of ICM-injected AAV to the brain since we expect this distribution route to work poorly for AAV5. Opposed to AAV5, AAV2-HBKO shows decreased uptake in peripheral organs compared to AAV9, which could also explain the success of FUS-MB delivery of this serotype to the brain after ICM administration if the primary route is distribution route #2(16, 24). AAVs have been optimized separately for increased brain transduction following ICM administration and FUS-MB-mediated delivery(25, 26). Assuming the importance of distribution route #2, the development of an optimal vector for ICM delivery combined with FUS-MB should focus both on increasing distribution following ICM-injection as well as decreasing uptake in peripheral organs.

## Conclusion

This study demonstrates that FUS-MB can expand delivery of ICM-administered AAV vectors to deep brain structures that are poorly reached through ICM injection alone. Further studies are warranted to elucidate the underlying mechanism of action, which is important for the design of future studies and clinical translation. ICM administration is becoming a promising route of administration for AAV-based gene therapies in clinical trials(12). The ability of FUS-MB to expand the footprint of AAV biodistribution to deep brain structures following ICM administration has the potential to significantly improve therapeutic efficacy in neurological diseases.

## Materials and methods

### Animals

Male Sprague Dawley rats (average 250 g) were ordered from Charles River either pre-cannulated with MR-compatible cannulas in the cisterna magna or without cannula. The cannula was flushed upon arrival with artificial cerebrospinal fluid (Bio-techne/Tocris, cat no 3525) in the volume indicated by the vendor as the void volume of the cannula. The experiments were conducted within 3 days of arrival of the animals to avoid clotting of the cannula prior to use. Animals were kept in a 12/12 light/dark cycle with food and water ad libitum and a temperature of 18° C - 22° C and humidity of 40%-60%. Animal work was performed according to the Canadian Council on Animals Care Policies & Guidelines and approved by the Sunnybrook Research Institute Animal Care Committee.

### Adeno-associated virus

AAV2-HBKO encoding GFP under a CAG promoter was produced as previously described using polyethyleneimine transfection of HEK293T cells(24, 27). All animals, irrespective of administration route, were injected with 5.4×10^11^ GC AAV2-HBKO in 45 μL sterile phosphate buffered saline (PBS) corresponding to a dose of 2.16×1012 GC/kg. Injection in intravenous catheters was followed by flushing with 0.2 mL 0.9% saline and injection in ICM cannulas was followed by flushing with artificial cerebrospinal fluid corresponding to the dead volume of the cannula as informed by the vendor.

### ICM injection of gadolinium

Gadolinium (Gadodiamide MRI contrast agent, Omniscan, GE Healthcare Canada, Mississauga, ON, Canada) was injected in the ICM cannula followed by flushing with artificial cerebrospinal fluid corresponding to the dead volume in the cannula. To mimic the injections of AAV2-HBKO, gadolinium was injected in a volume of 45 μL. T1-weighted MR images (500 ms TR, 6 ms TE, 256 × 256 matrix, 1.5 mm slice thickness) were acquired using a 7.0 T MRI instrument (BioSpin 7030; Bruker; Billerica, USA) 45, 55, 70 and 80 min following injection.

### FUS-MB treatment

For a detailed description of the FUS-MB treatment please see (28). Animals were anaesthetized with isoflurane and a tail-vein catheter was inserted followed by hair removal on the animal head to avoid trapping of air bubbles in the ultrasound gel. The animals were placed supine on an MR-compatible sled, and T2-weighted MR images (4000 ms TR, 70 ms TE, 256 × 256 matrix, 1.5 mm slice thickness) were acquired using a 7.0 T MRI instrument (BioSpin 7030; Bruker; Billerica, USA) and used for MR-guided targeting of the FUS. FUS was applied using a 0.58 MHz spherically focused transducer (75 mm outer diameter, 26 mm inner diameter, 60 mm radius of curvature) and an in-house manufactured system (prototype for LP100; FUS Instruments, Toronto, Canada). Definity microbubbles (0.2 mL/kg) were injected immediately upon FUS application of 10 ms bursts (burst repletion frequency 1Hz) at a fixed pressure of 0.32 MPa (measured in water) for 2 min for each target location. For intravenous injection of AAV2-HBKO, the injection was performed immediately after injection of microbubbles followed by intravenous administration of 0.2 mmol/kg gadolinium to visualize BBB permeability on T1-weighted MR images.

### Tissue collection

Four weeks following AAV delivery, animals were deeply anaesthetized using 75 mg/kg ketamine and 10 mg/kg xylazine followed by transcardial perfusion with 0.9% saline. Brains were collected; one hemisphere was post-fixed for 16 h in 4% paraformaldehyde in 0.1 M PO_4_, followed by transfer to 30% sucrose, and the other hemisphere was dissected in brain regions and flash frozen on dry ice. Organs and spines were collected and post-fixed for 16 h in 4% paraformaldehyde in 0.1 M PO_4_ followed by transfer to PBS.

### Immunohistochemistry and imaging

Brains were sectioned horizontally into 40 μm thick free-floating sections on a sliding microtome. Sections were washed three times for 10 minutes in PBS followed by antigen retrieval in 10 mM Tris base with 0.05% (v/v) Tween 20 and 1 mM EDTA (pH 9) at 70° C for 1 h. After allowing the sections to cool, washing in PBS wash repeated, followed by incubation for 2 h in blocking buffer (PBS with 0.3% Triton X-100, 3% (w/v) bovine serum albumin, and 10% (v/v) donkey serum). Sections were incubated with primary antibodies in blocking buffer overnight followed by washing in PBS and incubation overnight with secondary antibodies in blocking buffer. Staining with DAPI was performed by incubation for 10 min in PBS followed by washing in PBS, mounting on glass slides with polyvinyl alcohol medium and DABCO (Millipore, cat no 10981), and covering with glass coverslips. Primary antibodies include chicken anti-GFP 1:1000 (Abcam, ab13970) and guineapig anti-NeuN 1:500 (Millipore, ABN90). Secondary antibodies were purchased from Jacksom ImmunoResearch and diluted 1:400.

Whole section images were acquired with a 10x objective using a Zeiss Axio Scan.Z1 slide scanner. Images used for quantifications were acquired with a 20x objective using a Leica Stellaris with white light laser.

### RNA extraction and qPCR

Brain tissue was homogenized in 1 mL Trizol (Thermo Fisher, cat no 15596018) using a bead homogenizer followed by addition of 200 μL chloroform and 15 min centrifugation at 12,000xg. The supernatant was mixed 1:1 with 70% ethanol and RNA was extracted using PureLink RNA Mini Kit (Thermo Fisher cat no 12183018A) according to manufacturer’s protocol. RNA concentrations were measured using a Nanodrop 2000 (Thermo Fisher). One μg RNA was used for cDNA preparation using High-Capacity cDNA Reverse Transcription Kit (Thermo Fisher, cat no 4368814) according to manufacturer’s protocol. cDNA was diluted 1:50 and 5 μL was added to each well of a 385-well plate together with 1 μL of each reverse and forward primers (from a 10 μM dilution), and 7 μL SYBR Green qPCR master mix (Thermo Fisher, cat no 4472908). The following primers were used: GFP forward primer 5’-ACTACAACAGCCACAACGTCTATATCA-3′, GFP reverse primer 5’-GGCGGATCTTGAAGTTCACC-3′, and as reference genes Hprt1 forward primer 5’-TCCTCAGACCGCTTTTCCCGC-3’, Hprt1 reverse primer 5’-TCATCATCACTAATCACGACGCTGG-3’, Pgk1 forward primer 5’-ATGCAAAGACTGGCCAAGCTAC-3’, Pgk1 reverse primer 5’-AGCCACAGCCTCAGCATATTTC-3’. Samples were run on a QuantStudio 6 Pro Real-Time PCR System (Applied Biosystem) and the program: hold 50° C for 2 min, 95° C for 10 min, 40 cycles of 95° C for 15 sec and 60° C for 1 min. Results were analyzed using the 2^-ΔΔC^_T_ method with the average of the reference genes.

### DNA extraction and quantification of GFP genome copies

DNA was extracted from organs using QIAamp DNA FFPE Tissue Kit (Qiagen, cat no 56404) and from blood samples using DNeasy Blood and Tissue Kit (Qiagen, cat no 69504) according to manufacturer’s protocol except for the final elution step being done with a low TE buffer instead of the elution buffer provided in the kit. Digital droplet PCR analysis was conducted as previously described in detail(16). GFP primers were 50-ACT ACA ACA GCC ACA ACG TCT ATA TCA-30 and reverse primer 50-GGCGGATCTTGAAGTTCACC-30 (Invitrogen) and the probe was 50-6-FAM-CCG ACA AGC-ZENAGA AGA ACG GCA TCA-Iowa Black FQ-30 (Integrated DNA Technologies, Coralville, IA, USA). *Rpp30* reference gene primers and probe were obtained as a ddPCR Copy Number Assay (Bio-Rad, Part number 10042961, Assay ID dRnoCNS421683336).

### Spine analysis

Rat spines were excised by disconnecting the ribs and surrounding tissue. Subsequently, the spines were post-fixed overnight in a solution of 4% paraformaldehyde in 0.1 M PO_4_. The spine underwent a 15-min wash in PBS and was then incubated for approximately 30 days in 10% EDTA in PBS at 37° C, pH 7.5 and on a rotatory shaker. The EDTA solution was changed twice daily, and the bone was regularly inspected until it reached the necessary softness for sectioning. After another 15-min PBS wash, the spine was post-fixed in 4% paraformaldehyde for 1 hour at room temperature, followed by 15-min PBS wash. Spines were incubated in progressively increasing concentrations of sucrose in 0.1 M PO_4_, starting from 10% and incrementally reaching 20% and 30% sucrose daily, all at 4° C, shaking. Subsequently, a section from the thoracic part of the spines were embedded in an 8×8 mm tissue embedding mold (Electron microscopy sciences, 70180) by freezing in Tissue-Tek OCT (Sakura, Torrance, USA) and sectioned into 12-μm-thick sections using a Leica CM3050 S cryostat. Sections were mounted onto Apex Superior Adhesive slides (Leica, 3800080E-144) and stored at -80^0^C until staining. The spine sections were equilibrated to room temperature and washed 3 times 10 min in PBS, followed by incubation for 2 hours in blocking solution (PBS with 0.3% Triton X-100, 3% w/v bovine serum albumin, and 10% v/v donkey serum) in a humidify chamber. Sections were incubated overnight with primary antibodies in blocking solution at 4° C followed by three 10-min washes in PBS at room temperature. Subsequently, sections were incubated overnight at 4° C with secondary antibodies in blocking solution, followed by incubation in PBS with DAPI (1:10,000) (Sigma, D9542) for 10 min. Tissues were washed twice for 10 min in PBS at room temperature and ones for 10 min in 0.1 M PO_4_ before adding polyvinyl alcohol medium and DABCO (Millipore, 10,981), and covered with a glass coverslip. All solutions were directly applied to the mounted tissue. To prevent solutions from sliding off the slides a hydrophobic pen was utilized to outline the tissue’s edges, ensuring the localization of reagents on tissue specimens.

### Statistics

Normal distribution of qPCR data was determined using Shapiro-Wilk test. Non-transformed data from all brain regions was not normality distributed. Therefore, statistical analysis of non-transformed data in Supplementary Figure 1 was done using a non-parametric Kruskal-Wallis test with post hoc Dunn’s test. Log10 transformation resulted in normal distribution of data from the striatum, thalamus, cerebellum, and cortex 3, which was analyzed by one-way ANOVA and post hoc Tukey’s test. Log10 transformation of results from the hippocampus, brainstem, midbrain, cortex1, and cortex2 did not result in normal distribution, and statistical analysis of qPCR data from these brain regions was therefore done using a non-parametric Kruskal-Wallis test with post hoc Dunn’s test. All bars represent mean +/- standard deviation, ****p < 0.0001, ***p < 0.001, **p < 0.01, *p < 0.05. N numbers are indicated in each figure legend.

## Acknowledgements

The authors would like to thank Miguel Novelo, MSc, Nathalie Vacaresse, PhD, Melissa Rahmati, and Andrew Nicholson for their assistance with animal care, tissue collection and processing, and administrative tasks related to this study. The authors thank the Preclinical Research facility at Sunnybrook Research Institute for assistance with animal preparation for FUS-MB treatments. We are grateful to the Microscopy and Imaging Laboratory at the University of Toronto for guidance and access to the Zeiss Z1 slidescanner. We thank Dr. Yutaka Amemiya at the Genomics Core Facility at Sunnybrook Research Institute for assistance with ddPCR analysis and histotechnologist Petia Stefanova, MSc, for sectioning spines at the histology facility at Sunnybrook Research Institute. The authors would like to thank the GMU Vector Core at Sanofi for AAV production.

