## Supplementary figure 1 for "Focused ultrasound increases gene delivery to deep brain structure following the administration of a recombinant adeno-associated virus in the cerebrospinal fluid"

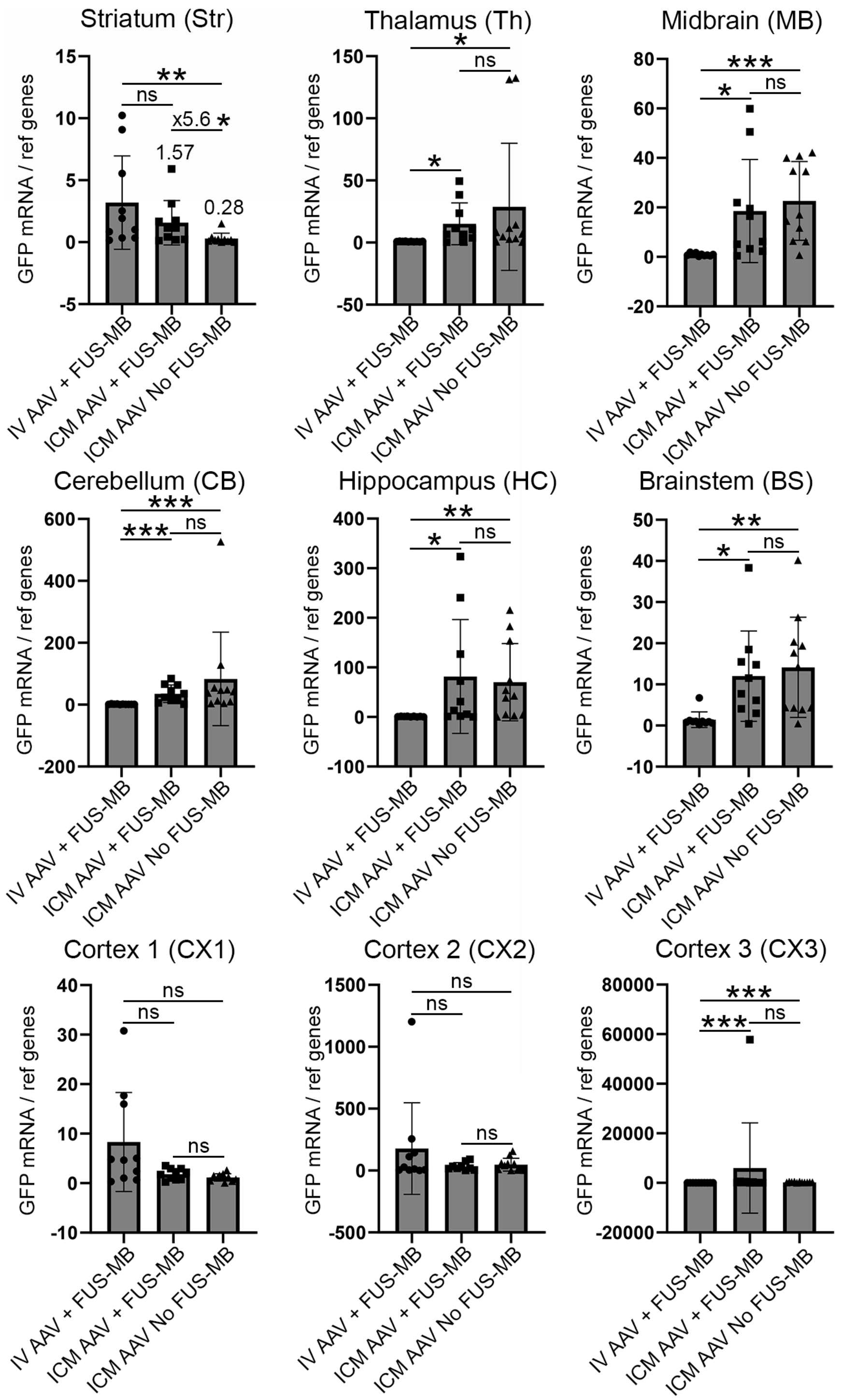


**Supplemental Figure 1 FUS-mediated delivery of ICM administered AAV to the striatum without log10 transformation**

Non-transformed data from quantification of GFP mRNA showed a 5.6 times improvement in GFP mRNA expression in the striatum in animals injected ICM with AAV and treated with FUS-MB compared to without FUS-MB. Statistical analysis was performed using non-parametric Kruskal-Wallis test and post hoc Dunn’s test because the data was not normally distributed. ***p < 0.001, **p < 0.01, *p < 0.05, n= 10-11/group.
